# Adherence rate to iron folic acid supplementation among pregnant women

**DOI:** 10.1101/535534

**Authors:** Wondwossen Niguse

## Abstract

**Introduction:** Globally 41.8% of pregnant women are anemic with the highest proportion affected in developing countries. Nationally, only 0.4% of the pregnant women take Iron supplements more than 90 days of the recommended 180 days. In Oromiya region 75.3% of pregnant women do not take any iron tablets or syrup during their last pregnancy, while 10.8% take for less than 60 days, 0.4% took for 60-89 days and only 0.3% took for 90 days or more.

**Objective:** To assess the adherence rate to iron and folic acid supplements among pregnant women attending antenatal clinics in Asella Town, south east Ethiopia

**Method and materials:** Institution based cross-sectional quantitative study design was conducted in Asella town from September 2015 to June 2016. A purposive sampling technique used to select the health institution. There are six health institutions selected for this study. The sample size 317 was selected with systematic random sampling method. Ten percent of pre-test was conducted in one institution which is not included in data collection. Data were collected using structured pre-tested questionnaire. Before data collection verbal consent was obtained. The collected data were analyzed using Epi-data and SPSS version 22.00 packages.

**Result:** The study revealed that Out of 317 pregnant women 296 (93%) responded to the questioner. The study revealed that 177 (59.8%) of pregnant mothers in the town adhered to the iron/folic acid supplement.

**Conclusion and recommendation:** Adherence of iron/folic acid supplementation found in this study is very low. Consequently, maternal education, adequate supplement supply to the health facility, early starting antenatal visit, and health education on duration of supplementation

## Introduction

Iron deficiency anemia is the most common nutritional disorder affecting two billion people worldwide (1). Based on evidence from iron supplementation trials, it was estimated that, on average, 50% of anemia globally is caused by iron deficiency (2). Pregnant women are at especially high risk of iron deficiency and anemia because of significantly increased iron requirements during pregnancy. Iron supplementation has been a major strategy in low-income and middle-income countries where micronutrient deficiencies are common to reduce iron deficiency anemia in pregnancy (2, 3).

Globally 41.8%, almost half of all pregnant women are anemic with the highest proportion affected in developing countries. The prevalence of anemia among pregnant women in developed country is 18% in average, which is significantly lower than the average 56% in developing countries. The actual prevalence of anemia in pregnant women in Africa and Asia is estimated to be 57.1% and 48.2%, while that of America and Europe is 24.1% and 25.1% respectively (4, 5).

Currently seventeen percent of Ethiopian women age 15-49 are anemic with the highest proportion of pregnant women (22%) than breast feeding (19 %) and neither pregnant nor breastfeeding women (15 %). Anemia prevalence also varies from urban and rural residence; a higher proportion of women in rural areas are anemic (18 %) than those in urban areas (11 %) (6).

The 2011 EDHS revealed that maternal nutritional status is poor in many respects in Ethiopia. Out of 17% of anemic women, 13% of them have mild anemia where hemoglobin level range between 10 g/dl and 10.9 g/dl, 3% having moderate anemia where Hgb level range between 7 g/dl and 9.9 g/dl, and 1% having severe anemia where Hgb level is <7 g/dl. (6,7,8).

Ethiopia, like most sub-Saharan Africa countries, has a national policy to prevent and treat anemia in pregnancy. This includes the provision of ferrous sulfate and folic acid to all pregnant women. The recommended dose in Ethiopia is 300 to 325 mg (milligrams) of ferrous sulfate and 400μg of folic acid once a day taken by mouth for 180 days of prenatal period, preferably with a meal. This dosage is usually supplied in a single combined iron and folic acid tablet (9).

## Methods and materials

### Study area

The study was conducted in Asella town south east Ethiopia. It is located in the Arsi Zone, Oromia Region about 175 kilometers from Addis Ababa. The current total population for Asella town is reported as 93,729, of whom 47,801(51%) were men and 45,927(49%) were women. Out of the total population 20,714(22.1%) of them were in reproductive age group. The majority of the inhabitants said they practiced Ethiopian_Orthodox Christianity, with 67.43% of the population reporting they observed this belief, while 22.65% of the population were Muslim, and 8.75% of the population were Protestant.

Azalea town comprises governmental (teaching and referral hospital and two health centers), nongovernmental (13 medium clinic, one hospital, one specialty MCH center and one higher clinic), nonprofit nongovernmental (FGA and Marie stops). Out of these 21 health institution eight of them comprise antenatal clinic out of which seven of them only provide a regular ANC check up.

### Study and Data collection period

The study period was from September 2015 to June 2016 and data collection period was April 1^st^ to 30^th^, 2016.

### Study design

Institutionally based cross-sectional quantitative study design was conducted, to determine the adherence rate to Iron-folate supplements among pregnant women in Asella town.

### Source population

The source population of the study was pregnant women attending ANC clinics in health institutions in Asella town.

### Study population

The study population was pregnant women attending ANC clinics in selected health institutions during the data collection period and that fulfill the inclusion criteria.

### Inclusion and exclusion criteria

#### Inclusion criteria

Pregnant women who had at least one ANC visit in health institution and supplemented with IFA tablets for at least one month before the date of interview.

#### Exclusion criteria

Pregnant women who come for the first antenatal visit, those who refuse to take the supplement, those mistakenly not provided the supplement, those who are unable to hear and/or speak and those who have mental disorder were excluded.

### Sample size determination

The sample size of this study was calculated by using the formula to estimate a single Population proportions

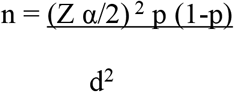

n = sample size,
Z α/2= significance level at α =0.05
P= established the prevalence from previous studies on the topic of interest (Adherence rate) in eight rural districts in Ethiopia (p=74.9%) (16).
d = margin of error of 0.05

Therefore, based on using the above single population proportion formula the sample size can be calculated as:

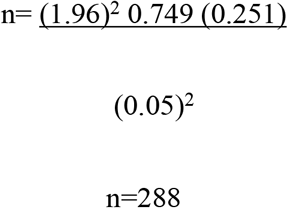

With the assumptions of the 95 % confidence interval, 10% non responsive rate the total sample size will be 317.

### Sampling procedure and technique

All health institutions (private and public) in Asella town were included in order to make the data representative. The health institution was selected with purposive sampling because the researcher was interested only on those institutions which provide regular ANC check up and a total sample size 317 pregnant women was selected using systematic random sampling as shown on the figure 1.

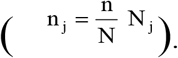

**Figure 1:**
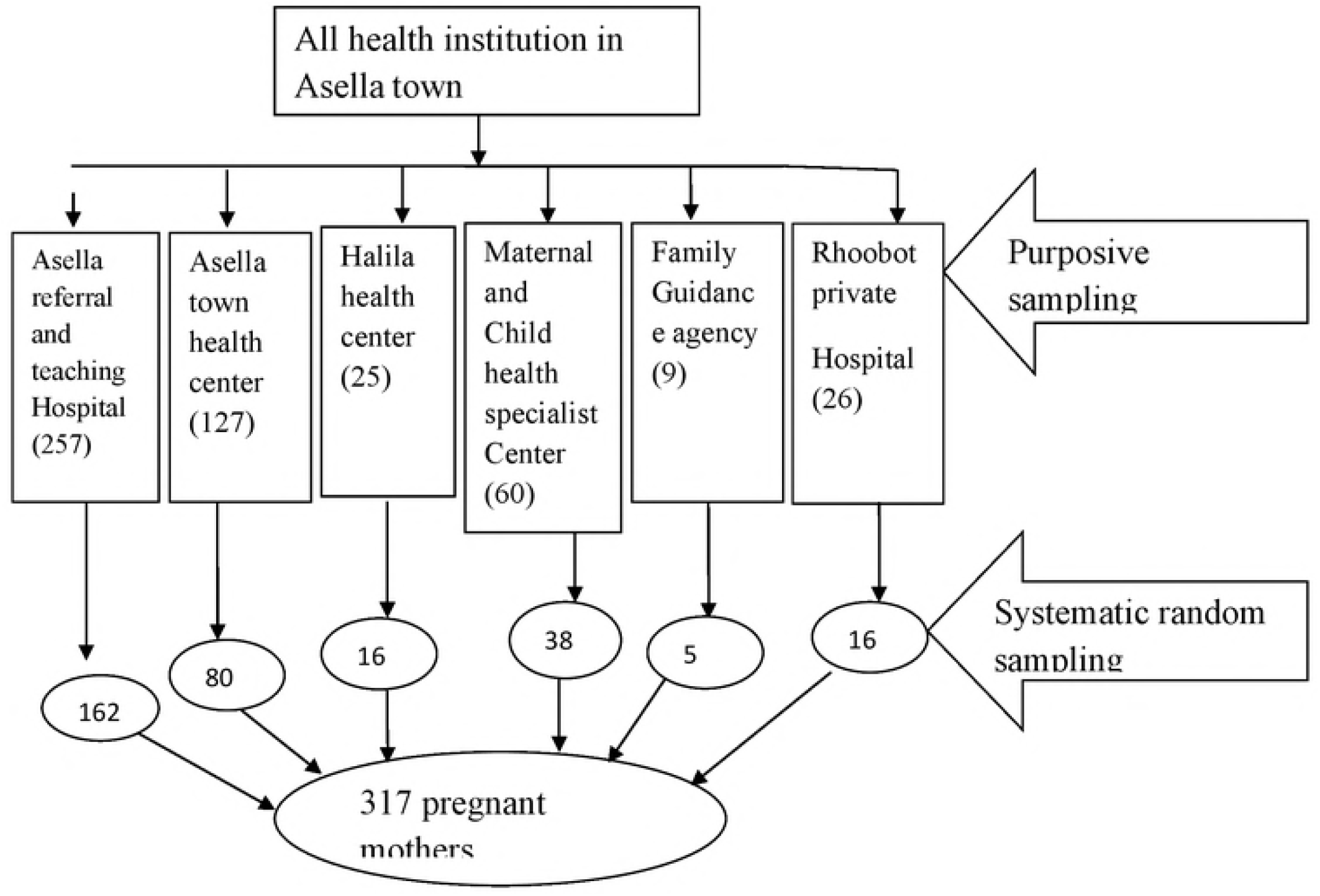
Schematic presentation of sampling procedure, Asella town, 2016. The total estimated number of pregnant women attending antenatal clinics in each antenatal institution for a single month was taken and proportional sample size was calculated for each institution so as to give the total sample size by using the following formula.

Where:

n_j_ = sample size of the j^th^ institution.
N_j_ = total population size of j^th^ institutions.
n = number of respondents to be selected from each institution.
N = Total number of pregnant women in selected institution (504).

|  |  |  |
| --- | --- | --- |
| For Asella referral hospital | $n_j = \frac{317 \times 257}{504}$ | = 162 |
| For Asella health center | $n_j = \frac{317 \times 127}{504}$ | = 80 |
| For Halila health center | $n_j = \frac{317 \times 25}{504}$ | = 16 |
| For specialist MCH center | $n_j = \frac{317 \times 60}{504}$ | = 38 |
| For FGA | $n_j = \frac{317 \times 9}{504}$ | = 5 |
| For Rhoboot hospital | $n_j = \frac{317 \times 26}{504}$ | = 16 |

317 pregnant mothers

### Dependent and independent variables

#### Dependent variable

Adherence to IFA supplement

#### Independent variables

1. Socio-demographic: - Age, marital status, socioeconomic status, family size, residence, maternal education and paternal Education
2. Obstetric factors: - Number of gravida, Parity, number of ANC visit, abortion, stillbirth, and ANC follow up started them.
3. Women Awareness: - Knowledge of anemia, knowledge about benefits of IFA and knowledge about duration of the supplementation

### Operational definitions

- **Adherence**: Pregnant women who had taken the combined iron/folic acid tablet at least 4 days a week considered to be adhered to the supplementation (6).
- **Non-adherence**: Pregnant women who had taken combined iron/folic tablet for less than 4 days a week considered to be not adhered to the supplementation (6).
- **Anemia**: Pregnant mother hemoglobin (Hgb) level in the blood is less than 11g/dl, which depicts decreased oxygen-carrying capacity for pregnant women.
- **Pregnant mothers Knowledge about anemia**: Those who score mean value (12.87) and above of correct response about cause, consequence, risk group and method of prevention of anemia are considered as they are knowledgeable about anemia while those who score less than the mean value (12.87) of correct response considered as they are not knowledgeable about anemia.
- **Pregnant mothers Knowledge about benefits of Iron/folic acid:** Those who score mean (1.24) value and above of correct response about benefits of iron/ folic acid considered as they are knowledgeable about benefits of iron, folic acid while those who score less than the mean value (1.24) of correct response considered as they are not knowledgeable about benefits of iron folic acid.
- **Antenatal care clinic (ANC):** A section in the health institution where a pregnant woman receives regular checkups, nutritional supplements (iron/foliate) and medical and nutritional information throughout her pregnancy.
- **Iron deficiency (ID):** A situation in which iron level found in the body of pregnant women recorded less than 3g.
- **Antenatal visits:** The pregnant woman optimally begins within 16 weeks of gestation and has four consecutive follow up at least each visit one month apart throughout the antepartum period.
- **Iron/folic acid supplement:** Additional combined nutrients to be taken by pregnant women in order to make up for iron/folic acid deficiency

### Data collection instrument and data collection procedure

Data collection instruments had closed and open ended questions and it includes six sections which are demographic, obstetric history, knowledge on anemia and folic acid, adherence to iron and folic acid and health care system factors. The questionnaire was adapted from a Kenyan study by Lynette awoke ding 2013 (36). Data was collected with structured pre-tested questionnaire using the interview. The questionnaire was prepared in English and translated into Afan Oromo then back to English, again, it translated into Amharic language and then back to English.

### Data quality assurance and management

Before the actual data collection, the questionnaire was pretested on 10% of similar population in Asella medium clinic to check the consistency and reliability. Data collector and supervisor were trained prior to conduct the data collection. After pretest some correction made to the questioner. Five diploma Nurses were recruited as a data collector and two BSc nurses were assigned for a supervisor, they were checking the data every day after data collection for the completeness of the questionnaire. Training was given to data collectors and supervisors for two days on purpose, of the study, details of the questionnaire, data collection procedure and filling the questioner.

### Data processing and analysis

The collected data were cleaned and checked for completeness; it was entered, compiled and analyzed with Epi data and SPSS version 22.00 packages. A univariate, was done using frequencies, to show adherence rate of iron and folic acid supplementation. Statistical significance declared at p-value less than 0.05.

### Ethical consideration

Ethical clearance was obtained from department of nursing and midwifery institutional ethical committee, school of allied health, College of Health Science. From the Department of Nursing and Midwifery the permission letter was written to Oromia regional health bureau to conduct the study and, then the permission letter was obtained from Oromia regional health bureau for the different study area to conduct the study. Finally informed verbal consent was obtained from each respondent. Information sheet and information was provided to the study participants about objective of the study. The response confidentiality was maintained.

### Dissemination and utilization of results

The result of the study will be disseminated to the Addis Ababa University College of health science school of allied health science department of nursing and midwifery, Addis Ababa University library, for each selected hospitals, MOH and for other concerned bodies through presentation, hard and soft copy. An attempt will be made for publication of the research on reputable Journal.

## Result

This chapter presents the study results including demographic and socioeconomic characteristics, factors hindering and those associated with adherence to iron/folate supplements, knowledge on anemia and iron/folic acid and frequency of ANC visits.

A total of 317 pregnant women attending antenatal care in governmental and private clinic in Asella town were included. Out of 317 pregnant women 296 (93%) responded to the questioner. 178 (60.1%) were from Asella town and 118 (39.1%) from the periurban area of the town. The reason for non-response to the survey was 24 pregnant mothers refused to be interviewed.

The age of the respondents ranges from 15-49 years. The mean age of the pregnant women was 27.7 and 27.2 years in rural and urban respectively. The majority of the respondents from urban and rural were married women [80.3%)] and [59.3%] respectively, and some of the women in the urban and rural area where single [15.2%] and [28.8%] respectively. While [4.5%] in urban and [10.2%] from rural were divorced.

Majority 47.8% and 22.5% of women from the urban area of the town had secondary and higher education respectively, while very few 11.9% and 2.5% of women from the rural area had secondary and higher education respectively. 39% and 2.8% of the respondents are illiterate from rural and urban women respectively. 44.4% and 10.2% of the husbands of the respondent were literate in the urban and rural area of the town respectively. Most of the husbands of the rural respondent [11.9%] were illiterate and 1.7% husbands of the respondent in the urban area were illiterate.

Selected socio-demographic variables of the study population are summarized in Table 1.

**Table 1:** Socio-demographic and economic characteristics of the study population, Asella town, 2016.

| **Variable** | **Urban**<br>**N=178 n (%)** | **Rural**<br>**N=118n (%)** |
| --- | --- | --- |
| **Age in years** |  |  |
| 15-19 | 8(4.7%) | 6(5.1%) |
| 20-29 | 119(66.9%) | 68(57.6%) |
| 30-39 | 50(28.1%) | 43(36.4%) |
| 40-49 | 1(0.6%) | 1(0.8%) |
| **Marital Status** |  |  |
| Single | 27(15.2%) | 34(28.8%) |
| Married | 143(80.3%) | 70(59.3%) |
| Divorced | 8(4.5%) | 12(10.2%) |
| Widowed | 0(0.0%) | 2(1.7%) |
| **Total number of Family size** |  |  |
| Below mean (2.7) | 145(81.5%) | 67(56.8%) |
| Above mean (2.7) | 33(18.5%) | 51(43.2%) |
| **Educational status of mother** |  |  |
| Can't read and write | 5(2.8%) | 46(39%) |
| Can read and write | 7(3.9%) | 36(30.5%) |
| Primary (grade 1-8) | 41(23%) | 19(16.1%) |
| Secondary (9-12) | 85(47.8%) | 14(11.9%) |
| Above secondary | 40(22.5%) | 3(2.5%) |
| **Educationl status of husband** |  |  |
| Can't read and write | 3(1.7%) | 14(11.9%) |
| Can read and write | 5(2.8%) | 37(31.4%) |
| Primary (grade 1-8) | 21(11.8%) | 33(28%) |
| Secondary (9-12) | 70(39.3%) | 22(18.6%) |
| Above secondary | 79(44.4%) | 12(10.2%) |
| **Continued table** |  |  |
| **Monthly income** |  |  |
| 151-650 | 9(5.1%) | 16(13.6%) |
| 651-1400 | 15(8.4%) | 41(34.7%) |
| 1401-2350 | 53(29.8%) | 22(18.6%) |
| **2351-3550** | 46(25.8%) | 19(16.1%) |
| **3551-5000** | 36(20.2%) | 18(15.3%) |
| **5001-10,000** | 19(10.7%) | 2(1.7%) |

A high percentage of the pregnant women interviewed were in their first trimester (35.5%),(33.4%) in their second trimester and (31.1%) were in the third trimester. Of the 296 pregnant women interviewed, 71.6% and 32.8% were multigravida and multiparous while 28.4% and 67.2% were primigravidae and primiparous (Table 2).

**Table 2:** Obstetric status of study women, Asella town, 2016.

| **Variable** | **Frequency(n)**<br>**(n=296)** | **Percent (%)** |
| --- | --- | --- |
| **Gravidity** |  |  |
| **Primigravida** | **84** | **28.4** |
| **Multigravida** | **212** | **71.6** |
| **Parity** |  |  |
| **Primiparous** | **199** | **67.2** |
| **multiparous** | **97** | **32.8** |
| **Number of ANC visits** |  |  |
| **2-3** | **260** | **87.8** |
| **>=4** | **36** | **12.2** |
| **ANC Starting Time** |  |  |
| **First trimester** | **105** | **35.5** |
| **Second trimester** | **99** | **33.4** |
| **Third trimester** | **92** | **31.1** |

When the respondents were asked if they had any history of birth complications such as stillbirths and abortion, 3.7% and 24.3% confirmed they had a history of stillbirth and abortion respectively while 96.3% and 75.7% did not have a history of asked complications. Majority 90.9% and 94.6% of women did not experience anemia neither in previous nor in current pregnancy (Figure 2).

**Fig 2:**
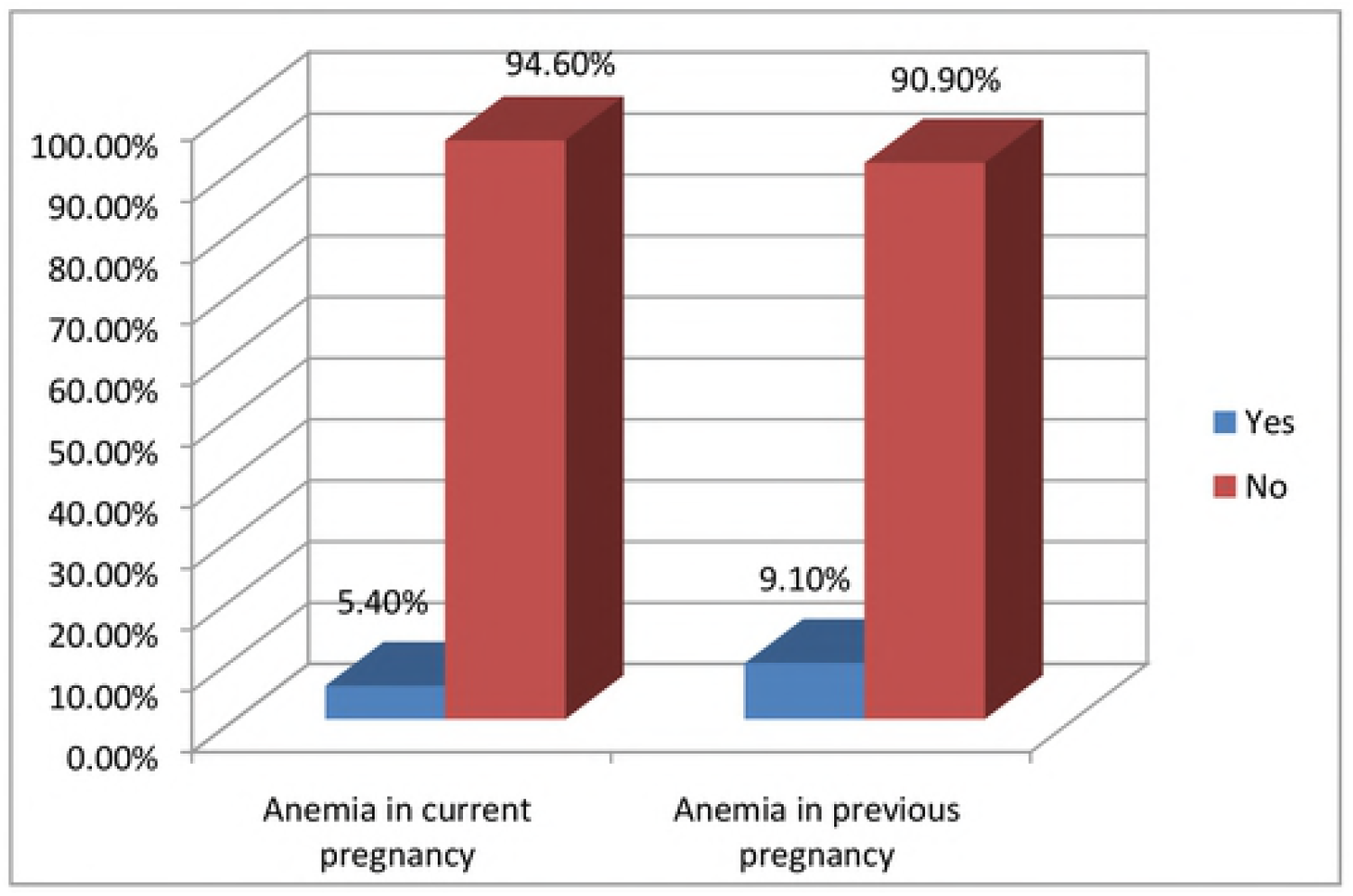
Previous and current history of anemia of the study women, Asella town, 2016. The great majority of interviews pregnant women [76%] and [50.7%] were knowledgeable about iron, folic acid and anemia respectively in giving the correct answer from listed correct and incorrect response to questions asked to type, duration, benefit and risk of IFA and cause, consequence, prevention of anemia and most susceptible group of people for anemia. While half, 49.3% of respondent mother were unable to give correct answer from listed correct and incorrect response to questions asked on anemia and Around one-quarter (24%) of the respondent mother fail to give correct answer from listed correct and incorrect response on iron and folic acid (Figure 3).

**Figure 3:**
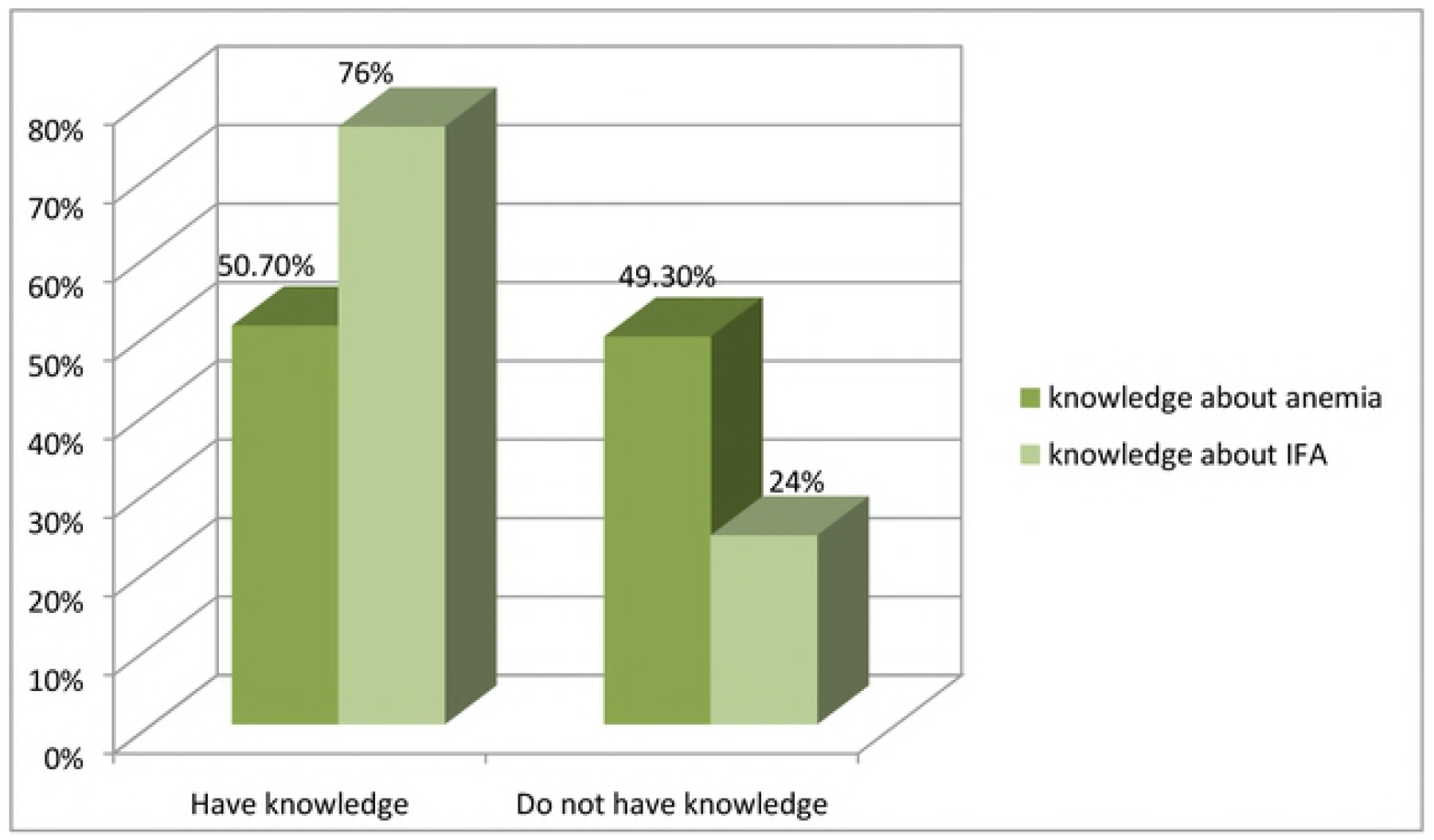
Respondents knowledge about anemia and iron/folic acid supplementation, Asella town, 2016. Health worker plays very great role in being Source of information for Most of pregnant women [60.6%] and [53.8%] about IFA and Anemia respectively, while school and friends were the least source of information for interviews women as shown in the figure 4.

**Figure 4:**
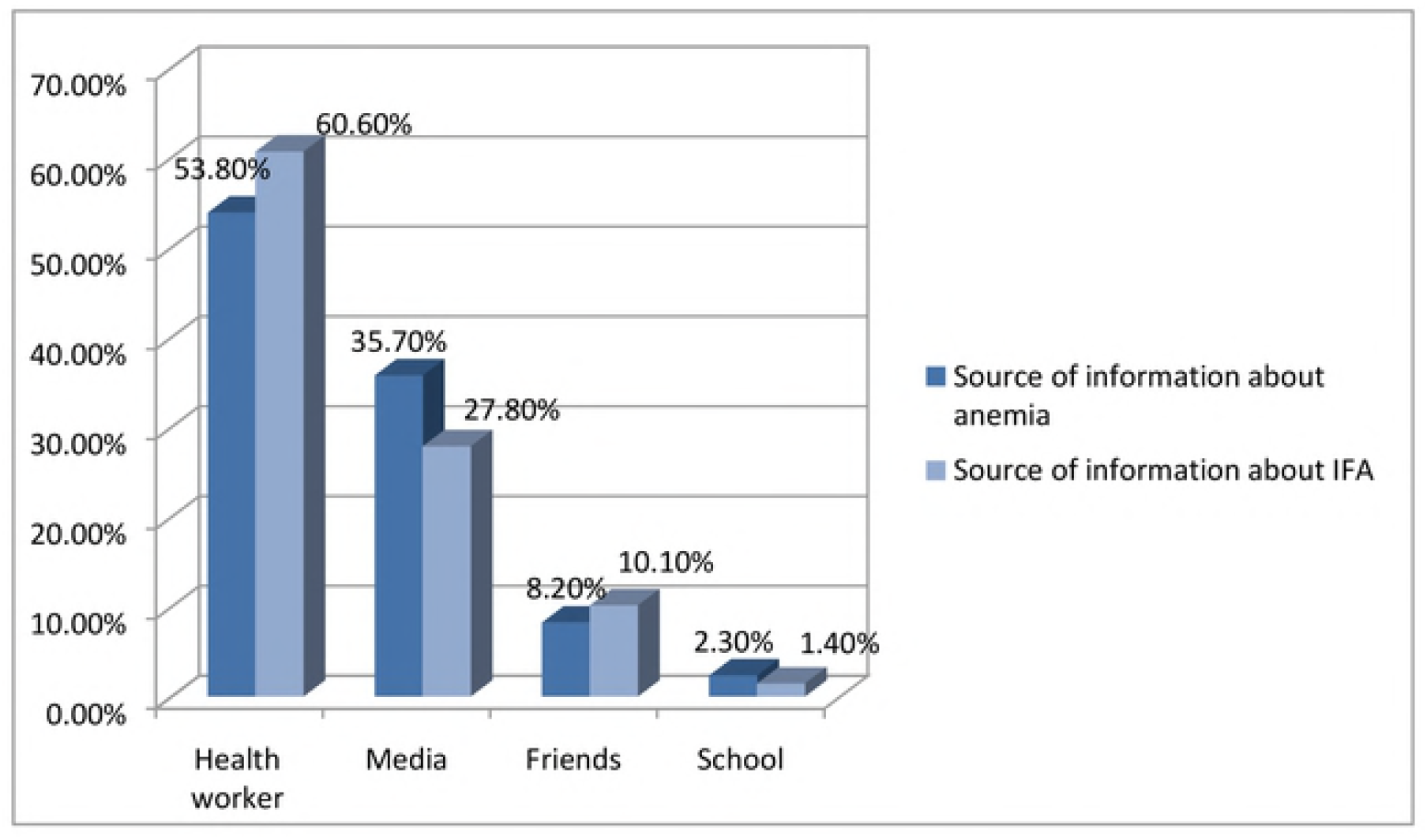
Source of information about anemia, Asella town, 2016. Adherence, which was considered as having taken a tablet of iron/folate supplements for four or more times in a week was observed by 177(59.8%) of the respondents, while 119(40.2%) had not adhered to the supplements (as shown in figure 5).

**Figure 5:**
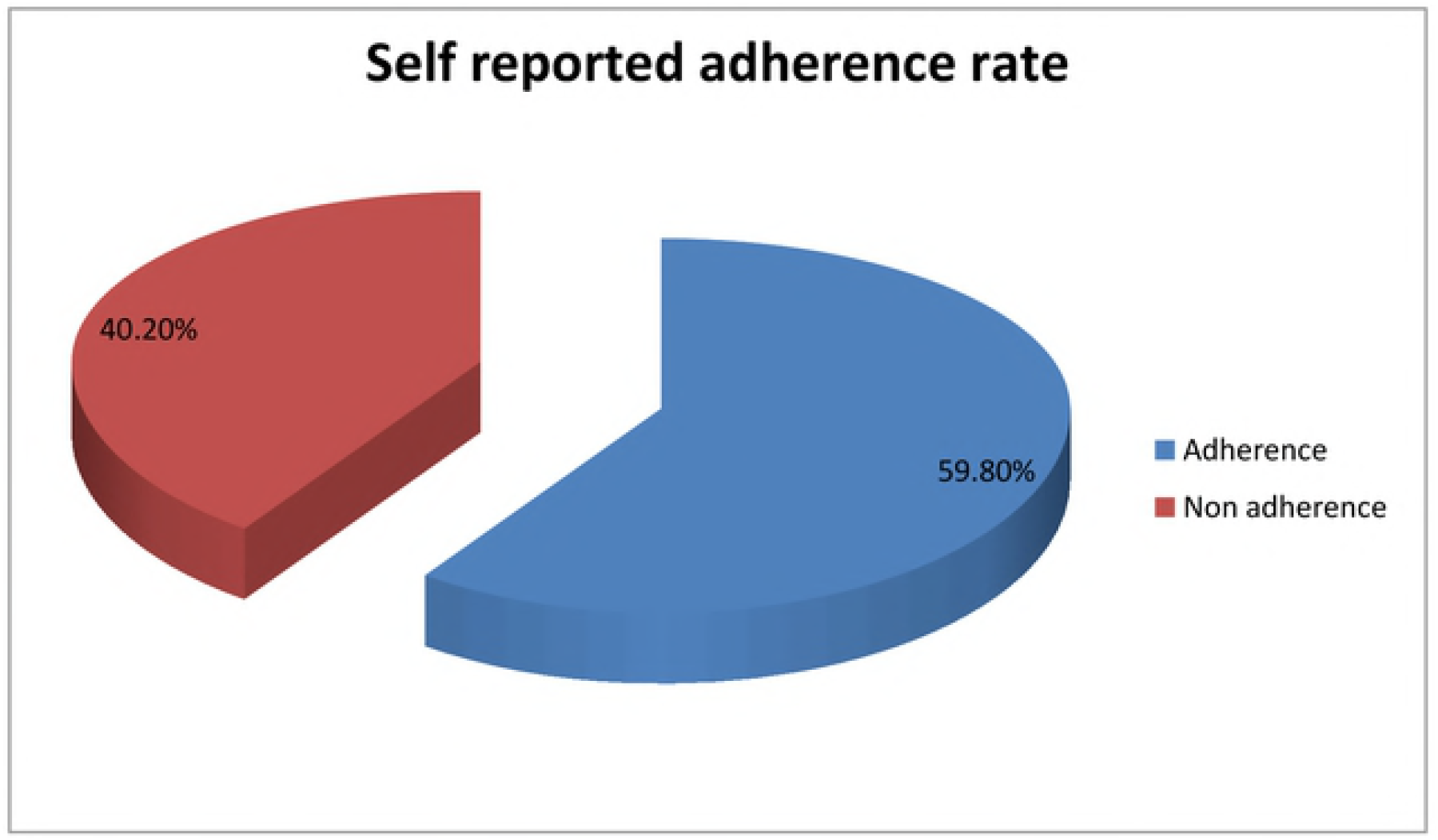
Self-reported adherences to the iron/foliate supplement, Asella town, 2016.

## Discussion

The adherence rate of iron and folic acid supplement found in this study is 59.8%, this result was consistent with studies done in the city of Mangalore, India, which was 62% and also consistent with the result found from cross-sectional study done in urban slums at Nagpur city, Maharashtra, India, which was 61.7% compliance. This consistency may be due to study were urban based and the pregnant women may get information from the health center, it may be due to health center accessibility (17, 30). Study done in Ethiopia, Oromia region, which shows adherence rate 0.3% is not consistent (6). In fact this study was done in urban areas and the women were probably more educated than those in rural areas could have contributed to the higher level than the regional level.

Iron/folic acid Adherence rate found in this study, 59.8% also higher than study done in Ethiopia, mecha district Amhara region result found was 20.4% (22). This difference in compliance may be due to the time gap, the culture of the people and different geographical location. And also cross-sectional study found in rural Kenya about adherence rate for optimum supplementation 90+ days was 18.3%. This difference is due to a study done in a rural set up, educational status, knowledge about supplement and access to the supplement are very low (19). In addition, result of adherence found in this study was lower than result found in eight rural districts in SNNP, Ethiopia, 2014 which was 74.9% average level of adherence rate of pregnant women in the area. This inconsistency may be due to cultural, geographical location and availability of drugs in the health center (16).

## Acknowledgment

First of all, I would like to express my deepest gratitude and appreciation to my advisor Rajalakshmi Murugan for her unreserved all rounded, support and enriching comment throughout the research thesis.

I would like to thank the department of nursing and midwifery, Addis Ababa University for giving this chance to prepare this research project. My appreciations also go to all staffs of school of nursing and midwifery for their unreserved support throughout the course and thesis works.

Finally, I would like to express gratitude for Asella Town Administrative Health office, the selected health institution in Asella town for providing the necessary information, the data collectors, supervisors and all participants who took part in this study.

